# Not all noise-reduction methods for fMRI preprocessing are created equal

**DOI:** 10.1101/2022.02.08.478718

**Authors:** M.E. Hoeppli, M.A. Garenfeld, C.K. Mortensen, H. Nahman-Averbuch, C.D. King, R.C. Coghill

## Abstract

Preprocessing fMRI data requires striking a fine balance between conserving signals of interest and removing noise. Typical steps of preprocessing include motion correction, slice timing correction, spatial smoothing, and high-pass filtering. However, these standard steps do not remove many sources of noise. Thus, noise-reduction techniques such as CompCor and FIX have been developed to further improve the ability to draw meaningful conclusions from the data. The ability of these techniques to minimize noise while conserving signals of interest has been tested almost exclusively in resting-state fMRI datasets and, only rarely, in task-related fMRI datasets. Application of noise-reduction techniques to task-related fMRI is particularly important given that such procedures have been shown to reduce false positive rates. However, little remains known about the impact of different noise reduction techniques on the retention of signal, particularly during tasks that may be associated with systemic physiological changes. In this paper, we compared two noise-reduction techniques, i.e. FIX and CompCor, in an fMRI dataset including noxious heat stimulation and non-noxious auditory stimulation.

Results show that preprocessing including FIX noise-reduction technique conserves significantly more signal than a preprocessing protocol including CompCor noise-reduction technique in both noxious heat and non-noxious auditory stimulations, while removing only slightly less noise. These results suggest that FIX might be the most appropriate technique to achieve the balance between conserving signals of interest and removing noise.

## Introduction

Functional MRI brain images, whether they result from resting-state or task-related sequences, are very noisy by nature (Fassbender et al., 2017; Network et al., 2013). The images need to be carefully preprocessed in order to filter out the noise while conserving the signal of interest (Fassbender et al., 2017; Johnstone et al., 2006; Lund et al., 2005; Oakes et al., 2005). Noise can emerge from different sources including motion, in particular head motion, MRI scanner artifacts, or physiological noise arising from normal cardio-respiratory activity (Brooks et al., 2013; Fassbender et al., 2017; Lund et al., 2005; Network et al., 2013; Yoshikawa et al., 2020). In the case of task-related MRI sequences, the task itself can add noise to the data (Fassbender et al., 2017). This additional noise can be due to the inherent nature of the task participants are asked to perform, such as any movement participants would have to complete in the task, or to any physical or physiological reaction to the performed task, in addition to noise introduced by equipment used during the task (Brooks et al., 2013; Epstein et al., 2007; Kasper et al., 2017; Yoshikawa et al., 2020).

Standard steps to preprocess the data typically include motion correction, slice timing correction, spatial smoothing, and high-pass filtering. Motion correction is the process by which the head movements are corrected by realigning volumes across the scan (Jenkinson, 2002; Poldrack et al., 2009). Slice timing correction controls for the differences of acquisition time between slices of each volume (Poldrack et al., 2009). Spatial smoothing consists in the application of a filter to remove high-frequency spatial noise, allowing better signal detection in large brain areas (Poldrack et al., 2009). Finally, high-pass filtering removes the low-frequency signals considered noise of physiological or scanner origin (Poldrack et al., 2009). In some cases, intensity normalization is also applied to the data, which results in a constant mean volume intensity over time (Coghill et al., 1994).

Numerous noise-reduction techniques have been developed to further clean the data, including the FMRIB’s ICA-based Xnoiseifier (FIX) and the Component-based correction method (CompCor) techniques, which are the two techniques compared in this manuscript. The first technique, FIX, was developed in the context of the Human Brain Connectome Project to filter out noise of various origins from resting-state fMRI data (Salimi-Khorshidi et al., 2014). FIX relies on the usage of a classifier to identify Independent Component Analysis (ICA) components as noise or signal (Griffanti et al., 2014; Salimi-Khorshidi et al., 2014). Usage of a classifier allows to reduce the workload compared to manual labelling and cleaning of the data, while preserving accuracy in the classification of the components. Although trained classifiers from the Human Connectome Project are available, it is recommended that the classifier is hand-trained on one’s dataset to further ensure accuracy of the classification (Salimi-Khorshidi et al., 2014). Recent studies (Blasi et al., 2020; Mayer et al., 2019) highlighted the successful use of FIX to denoise task-related fMRI data.

The second noise-reduction technique, CompCor, is widely used to denoise resting-state and task-related fMRI data. Noise is identified by performing a Principal Component Analysis (PCA) on signals from the white matter and cerebrospinal fluid (CSF). These signals are then regressed out of the data by performing a General Linear Model (GLM) analysis (Behzadi et al., 2007). Its efficiency has been demonstrated in perfusion and Blood-Oxygen Level-Dependent fMRI sequences (BOLD) (Behzadi et al., 2007).

Although the efficiency of these noise-reduction techniques has been demonstrated on task-related and resting-state fMRI data, the tasks included in these tests were often associated with language or visual skills. These tasks have the advantage of producing clear, strong signals in well-known areas, making them ideal to test noise-reduction techniques. In addition, these tasks do not typically induce much additional noise.

There is a need to test further the efficiency of these techniques in tasks that induce global cerebral blood flow changes (Coghill et al., 1998; Zeidan et al., 2015) and/or have substantial physiological responses or movements associated with them (Brooks et al., 2013; Perlaki et al., 2015; Tousignant-Laflamme et al., 2005), such as experimentally induced pain, alterations in breathing, or fear. For such tasks, it is essential to ensure that the noise-reduction technique removes the noise without removing the signal of interest, even if this signal might be present in areas typically associated with noise, such as the white matter.

This paper aims to compare the performance of a FIX noise-reduction technique with a CompCor noise-reduction technique and define the most appropriate technique to conserve signals of tasks associated with substantial changes in the white matter and/or in global cerebral blood flow(Coghill et al., 1998; Zeidan et al., 2015). A standard preprocessing, which does not include any noise-reduction technique, is used as the baseline to compare the two noise-reduction techniques. Because FIX is designed to detect spatial and temporal features of noise specific to one’s dataset, we hypothesize that the FIX noise-reduction technique would be more sensitive to conserving signal of interest while removing the noise than the CompCor noise-reduction technique.

## Methods

### Participants

143 healthy individuals (age range: 14 to 44 years old) were enrolled in a neuroimaging study investigating the individual experience of pain. Data from 34 of these participants (Table 1; age: 28 ± 6.1, mean ± SD), which were not used in any prior analyses (Hoeppli et al., 2020), were used to train an automated classifier FSL FIX (FMRIB’s ICA-based Xnoiseifier, FSL, Oxford, UK) (Griffanti et al., 2014; Salimi-Khorshidi et al., 2014) that was used to denoise the fMRI data. 4 of these 34 participants had subtle incidental findings reported by a radiologist, that due to their nature and localization should not affect the training of the FIX classifier. As a consequence, these 4 participants were not excluded from the classifier training. Out of the remaining 109 participants, 8 were excluded because of insufficient quality of the fMRI images or incidental findings of abnormalities on MRI. The remaining 101 healthy volunteers (Table 1; 43 males and 58 females, age: 28.5 ± 7.7, mean ± SD) were included in the analyses described in this manuscript.

**Table 1.** Demographics of the training and test datasets.

|  | **Training data**<br>**(N = 34)** | **Test data**<br>**(N = 101)** |
| --- | --- | --- |
| **Sex** |  |  |
| Females | 22 (64.7%) | 58 (57.4%) |
| Males | 12 (35.3%) | 43 (42.6%) |
| **Age** |  |  |
| Mean (SD) | 28 (6.1) | 28.5 (7.72) |

Participants and parents/legal guardians of minor participants gave their written informed consent and minors provided written assent in accordance with the institutional review board of Cincinnati Children’s Hospital Medical Center, which approved the study. Exclusion criteria included active neurological or psychiatric disorders that impacted the participant’s ability to perform the tasks requested, the presence or history of chronic pain, medications that could interfere with quantitative sensory testing (QST) or brain function, positive screen for recreational drugs, any serious pathology, substantial uncorrected visual deficit, and any MRI contraindication, such as any metallic implant or braces.

### General design

Participants in this study underwent two sessions: a QST session and an MRI session. The goal of the QST session was to familiarize participants with the stimuli and rating scale used in the MRI session. This session will not be further discussed in this manuscript, but a detailed description of this session can be found elsewhere (Hoeppli et al., 2020). The MRI session lasted approximately 90 minutes. During this session, participants experienced three types of stimuli: heat, cold, and auditory stimuli. After each stimulus, participants were asked to rate their pain intensity and unpleasantness sensations on two computerized visual analog scales. A complete description of the scale can be found in Hoeppli et al. (Hoeppli et al., 2020). Only the heat and auditory fMRI series will be detailed below; since analyses reported in this manuscript were performed on these stimulus modalities only. An in-depth description of the cold stimulus modality is available in Hoeppli & al. (Hoeppli et al., 2020). We chose to focus on the heat pain modality, as it is one of the stimulus modalities most frequently used to experimentally induce pain and is associated with a physiological response that might impact the MRI response. The auditory stimulus modality was chosen as a control condition for the heat modality since it is typically not associated with the same level of white matter response as noxious heat stimulation.

A parent or legal guardian accompanied minor participants during both sessions to confirm eligibility to participate in the study (QST session) and MRI compatibility (MRI session). Once eligibility was confirmed, parents were asked to step out to avoid influence on participants’ responses (McMurtry et al., 2010; Schinkel et al., 2017; Zohsel et al., 2006).

All participants were asked to turn their cell phones off during both sessions to avoid distractions and interruptions. During the MRI session, participants’ cell phones were kept secure with their other ferromagnetic belongings in lockers.

### MRI session

#### fMRI stimuli and Block-design fMRI series

##### Heat fMRI series

Participants completed three fMRI series heat stimuli followed by a 16-second rating period and a 22-second resting period. Participants received a total of 17 high intensity noxious stimuli (48°C) and 4 low intensity noxious stimuli (47°C). Two series included 6 high intensity and 1 low intensity noxious stimuli each, and one series included 5 high intensity and 2 low intensity noxious stimuli. Stimuli were delivered to the back of the lower left leg using a 16×16mm thermode provided by a Medoc pathway system (Medoc, Ramat Yishai, Israel). To limit sensitization or habituation to the stimuli, the position of the thermode was slightly moved on the participant’s calf between heat series. The repetition of the heat fMRI series was meant to increase statistical power at the individual level and decrease false positive rates. For each stimulus, the temperature increased from a baseline of 35°C at a rate of 6°C/s, plateaued for 10 seconds, and returned to baseline at a rate of 6°C/s.

Participants were instructed to rate their perceived pain intensity and unpleasantness after each stimulus. Between the presentation of the scales, a black screen was displayed.

##### Auditory fMRI stimuli

Participants completed one auditory fMRI series. This series included 5 high intensity non-noxious stimuli (90 dB) and 2 low intensity non-noxious stimuli (80 dB). The auditory stimuli were designed as 900Hz sawtooth waves with an overall time course similar to the heat stimuli. In the high intensity stimuli, the sound increased from silence to 90 dB in 2.2 seconds and plateaued for 10 seconds before returning to 0 dB at the same rate. In the low intensity stimuli, the sound increased to 80 dB in 2 seconds and plateaued for 10 seconds before returning to 0 dB at the same rate. After each auditory stimulus, participants were instructed to rate the intensity and unpleasantness of the sounds that they experienced.

#### MRI acquisition

During the MRI session, participants lay in a supine position in a Philips 3T Ingenia scanner with a 32-channel head coil. During this session, all participants always underwent a T1 structural scan first. They then completed three BOLD fMRI series of heat stimuli, one fMRI series of cold stimuli and one fMRI series of auditory stimuli. Resting BOLD and arterial spin label (ASL) series were acquired but are not reported here. The order of the BOLD and ASL series was counterbalanced between participants.

A radiologist inspected the structural images of the participants for incidental findings.

#### T1 structural scan

The multi-echo (4 echoes) T1-weighted series was acquired using the following parameters: repetition time (TR): 10msec; echo times (TE): 1.8, 3.8, 5.8, 7.8; flip angle: 8; FOV: 256 × 224 × 200 mm; voxel size: 1 × 1 × 1 mm; slice orientation: sagittal. The total duration of this scan was 4 minutes 42 seconds.

#### BOLD fMRI

Each functional image series consisted of 193 volumes acquired using the following parameters: TR: 2 sec; TE: 35msec; voxel size: 3 × 3 × 4 mm; FOV: 240 × 240 × 136 mm; slice orientation: transverse; slice order: ascending; dummy scans: 2. Each series lasted 6 minutes 26 seconds, after an 8-second pre-scan time.

##### Statistical analyses on fMRI data

All MRI data were first inspected for motion and scanner artifacts. They were then preprocessed and analyzed with FSL (FMRIB Software Library, version 6.0.1 Oxford, UK).

###### Data Preprocessing

Structural images were first corrected for bias using FMRIB’s Automated Segmentation Tool (FAST) {Zhang:2001ve}. Images were then brain extracted using the Brain Extraction Tool (BET) {Smith:2002ef} and normalized into standard space MNI-152 using FMRIB’s Linear Image Registration Tool (FLIRT) (Jenkinson, 2002; Jenkinson and Smith, 2001). Finally, images were segmented into the different tissue types and white matter and CSF were masked at a probability threshold of 0.95.

fMRI data from 135 participants were used in these analyses. 34 participants were included in the training dataset for a FIX classifier. Data from the remaining 101 participants were included in the test dataset to compare two preprocessing protocols with noise-reduction techniques, i.e. FIX and CompCor, to a standard preprocessing protocol for fMRI data without noise-reduction technique (Figure 1). In each preprocessing protocol, data were visually inspected after each preprocessing step to confirm that the correction was correctly applied to the data.

**Figure 1.**
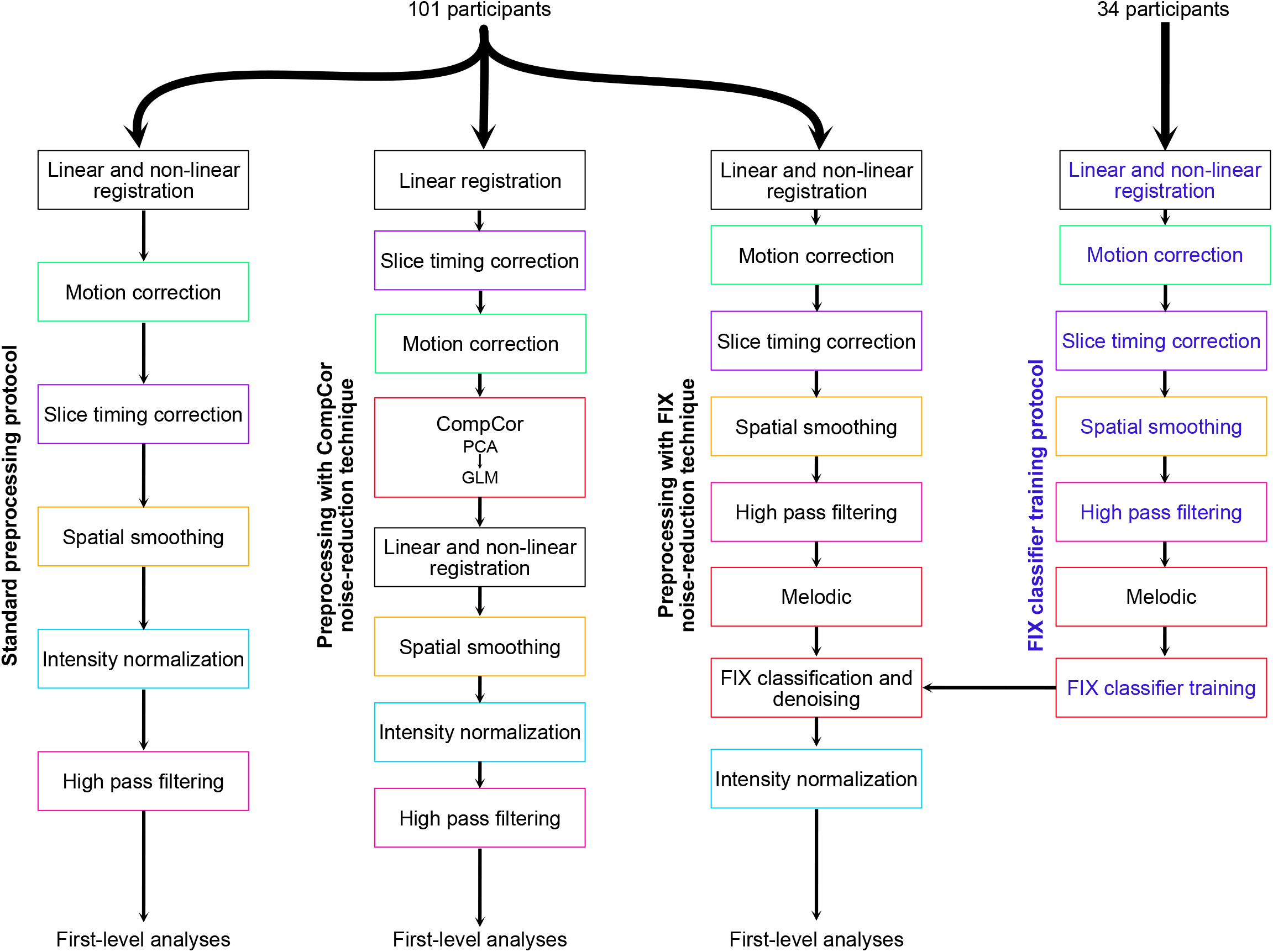
Flowchart of the three main preprocessing protocols, including standard preprocessing, preprocessing with CompCor noise-reduction technique, and preprocessing with FIX noise-reduction technique, and the FIX classifier training protocol. PCA: Principal Component Analysis; GLM: General Linear Model

#### Standard preprocessing protocol

The test data were preprocessed using a standard preprocessing protocol (Figure 1). This protocol included the following steps: motion correction using MCFLIRT (Motion Correction FMRIB’s Linear Registration Tool) (Jenkinson, 2002), slice timing correction, brain extraction (BET), spatial smoothing (FWHM: 5mm) with SUSAN (Smith and Brady, 1997), high pass filter (cutoff: 100 sec), and intensity normalization. No additional noise-reduction technique was used to further remove noise from the data.

#### Preprocessing protocol with FIX noise-reduction technique (FIX preprocessing)

##### Preprocessing of the training dataset

The training datasets were first registered to the structural scan and then to the standard space MNI-152 using FLIRT and FNIRT (FMRIB’s Non-linear Image Registration Tool) (Andersson et al., 2007; Jenkinson et al., 2012; Smith et al., 2004; Woolrich et al., 2009). These datasets were then preprocessed in the following steps: motion correction using MCFLIRT (Motion Correction FMRIB’s Linear Registration Tool) (Jenkinson, 2002), slice timing correction, brain extraction (BET), spatial smoothing (FWHM: 5mm) with SUSAN (Smith and Brady, 1997), and high pass filter (cutoff: 100s).

After these preprocessing steps and before training the FIX classifier, a Probabilistic Independent Component Analysis (PICA) using MELODIC (Multivariate Exploratory Linear Optimized Decomposition into Independent Components) (Beckmann and Smith, 2004) was performed to extract the components to classify.

To define the ideal number of components for our dataset to separate noise from the signal, MELODIC was performed using four predefined numbers of output independent components (15, 20, 25, and 30 components) on a random subsample of 10 participants with three fMRI heat series each, i.e. 30 datasets. It was determined that PICA performed using MELODIC and resulting in 25 output independent components (IC) was best at separating noise from signal in our data.

Thus, PICA performed using MELODIC and outputting 25 ICs was then performed on our complete training dataset. Components resulting from MELODIC were then manually categorized by three experts (MEH, MAG, and CKM) as signal, noise, or unknown when the source of the activation could not be defined unambiguously. Categories were defined based on the spatial map, the power spectrum, and the time series of the IC. All the categories were defined according to Salimi-Khorshidi et al. (Salimi-Khorshidi et al., 2014). ICs identified as noise were then subcategorized into “Unclassified noise”, “Movement”, “MRI”, “Respiratory”, “White matter”, “Susceptibility motion”, “Sagittal sinus”, and “Cardiac”. A single IC identified as noise could be labeled with multiple subcategories of noise.

Once consensus was achieved between the three experts, categorized ICs of the 102 datasets (34 participants x 3 fMRI heat series) were used to train a FIX classifier, creating a trained-weights file to be used in test datasets to automatically categorize ICA components (FMRIB’s ICA-based Xnoiseifier, FSL, Oxford, UK) (Griffanti et al., 2014; Salimi-Khorshidi et al., 2014). Finally, a leave-one-out test was performed to assess the efficiency of the classifier to correctly identify components as signal and noise and define the optimal distinction threshold for our data.

##### Preprocessing of the test datasets

Test datasets were preprocessed following the FIX preprocessing protocol (Figure 1). This protocol included the same steps as those described in the training data, i.e. motion correction, slice timing correction, BET, spatial smoothing (FWHM: 5mm), and high pass filter (cutoff: 100s). Next, PICA was performed on the preprocessed data using MELODIC and was set to output 25 ICs. Once ICs had been defined by MELODIC, the FIX classifier, which was trained on our training dataset, was applied to the ICs of the test datasets to categorize them into signal and noise and remove the components identified as noise. The threshold to separate noise and signal components was defined to preserve as much signal as possible. The same classifier was used for the preprocessing of heat and auditory fMRI series. Before applying it to the auditory fMRI series, the accuracy of the classifier to distinguish noise from signal of interest in these data was confirmed by performing a leave-one-out test. The test confirmed the accuracy of the classifier on the auditory data. Finally cleaned filtered images were corrected by intensity normalization.

#### Preprocessing protocol with CompCor noise-reduction technique (CompCor preprocessing)

The test dataset was preprocessed using the following protocol including a CompCor noise-reduction technique (Figure 1). Data underwent slice timing correction and motion correction before being linearly registered to the T1 image. A CompCor noise-reduction technique was then applied to the data. This technique included a principal component analysis (PCA) and General Linear Model (GLM) analysis (Behzadi et al., 2007). The PCA was performed on the activation time series extracted from the white matter and CSF to create nuisance regressors. These regressors were then used to create the design matrix for the GLM analysis and filter out noise. This technique is based on two assumptions (Behzadi et al., 2007): (1) The activation present in the white matter and CSF is only physiological noise and not signal of interest; (2) Noise in the grey matter is correlated to noise in the white matter and CSF and can be filtered out by regressing time series of noise in white matter and CSF. Denoised data were then registered linearly and non-linearly to native and standard space. Finally, the data underwent spatial smoothing, intensity normalization, and high-pass filtering to complete the preprocessing protocol.

#### Data Analyses

##### Overview

Five analyses were performed to evaluate the impact of the three preprocessing protocols on the signal-to-noise ratio qualitatively and quantitatively. Qualitative comparisons included block-design based analyses of the cerebral responses to high intensity heat stimulus and of the percentage of participants exhibiting the same brain activation (frequency analysis). T-tests using a block design were then performed to quantitatively compare brain activation differences between the preprocessing pipelines. These T-tests were repeated using a single epoch design to compare differences in activation associated with a single epoch, i.e. one noxious stimulus, between the preprocessing pipelines. T-tests using a block design were also replicated in the auditory modality to investigate the effect of the three preprocessing protocols on a task with a lower level of white matter response. Finally, contrast parameter estimates (copes) and variance of contrast parameter estimates (varcopes) were compared to investigate differences in the detected signal (copes) and noise (varcopes) between our preprocessing protocols. For all fMRI analyses, a clustering z threshold of 3.1 and a p threshold of 0.05 were used (Eklund et al., 2016).

##### Differences in brain activation associated with high intensity heat and auditory stimulation using a block design between the preprocessing protocols

To qualitatively compare the effect of the three preprocessing protocols on the data, GLM analyses were performed to define brain activation associated with high intensity heat and auditory stimulus.

First-level GLM analyses were performed on each individual fMRI heat and auditory series preprocessed with each protocol using FEAT (Woolrich et al., 2001). One block-design regressor of interest, i.e. high intensity stimuli, and two block-design regressors of no interest, i.e. low intensity stimuli and rating periods, were defined. For the heat paradigm, which included three series, individual second-level GLM analyses for each preprocessing protocol were performed on contrast parameter estimates (cope) images derived from the first-level heat GLM analyses using FEAT (Woolrich et al., 2004), allowing combination of the individual heat series using a fixed-effect statistical model. Finally, a group-level GLM was performed with FEAT (Woolrich et al., 2004) using cope images from the individual second-level heat analyses and the individual first-level auditory analyses using mixed-effect FLAME 1 and 2 statistical models.

To quantitatively identify the differential impact of the preprocessing protocols on brain activation associated with high intensity heat stimulus (48°C), three individual T-tests were then performed on the individual second-level copes using a fixed-effect statistical model. The first T-test compared brain activation following the FIX preprocessing protocol to the activation following the standard preprocessing protocol. The second T-test compared brain activation following the CompCor preprocessing protocol to the activation following the standard preprocessing protocol. The third T-test compared brain activation following the FIX preprocessing protocol to the activation following the CompCor preprocessing protocol. Finally, three group-level GLM analyses were performed on the individual differential copes using a mixed-effect FLAME 1 and 2 statistical model to investigate differential effects of the preprocessing protocols on brain activation associated with high intensity heat stimuli at a group level.

Similarly, the same three T-tests were performed on the individual first-level auditory copes, followed by similar group-level GLM analyses as described above to investigate the differential effects of the preprocessing protocols in a task without important associated white matter response.

##### Frequency of activation across participants

The frequency (percentage) of participants exhibiting activation in brain areas commonly activated in response to heat stimuli was calculated to assess differences in within-individual activation between the processing protocols.

Individual second-level block-design copes of the heat paradigm were concatenated, resulting in a 4D dataset using fslmerge. This dataset was then binarized and averaged across time using fslmaths. This allowed us to examine the percentage of participants showing activation in brain areas relevant to the used paradigm.

##### Differences in brain activation associated with a single high intensity heat stimulus using a single-epoch design between preprocessing protocols

To further evaluate the differential impact of the preprocessing protocols on brain activation associated with high intensity heat stimulus, first-level individual GLM analyses were repeated using a single-epoch design, where each epoch represented one stimulus of the first heat fMRI series. Single-epoch designs are more susceptible to noise than block designs (Koyama et al., 2003). Therefore, the impact of the preprocessing protocols should be more pronounced in a single-epoch design. The first stimulus of that series was defined as a regressor of interest, while the other 6 stimuli and rating period were defined as regressors of no interest. Two contrasts were defined as positive (contrast 1) and negative (contrast 2) activation associated with the first stimulus. Group-level GLM analyses were then performed for each protocol to identify positive and negative brain activation associated with that stimulus across participants. Three second-level individual T-tests, which included similar comparisons as the ones done on the main block-design analysis, were then performed on the individual first-level copes to statistically compare brain activation associated with a single high intensity heat stimulus between the preprocessing protocols. Finally, three group-level GLM analyses were performed on the individual differential copes using a mixed-effect FLAME 1 and 2 statistical model to investigate differential effects of the preprocessing protocols on brain activation associated with a single high intensity heat stimuli across participants.

##### Noise and signal detection following each preprocessing protocol

To further investigate the effect of each preprocessing protocol on the signal of interest and noise removal, we isolated the parameter estimates of individual second-level block-design copes and varcopes averaged within areas of interest, including ACC, S1, insula, and within areas considered as sources of noise, i.e. white matter and CSF. We then compared these estimates within areas and between preprocessing protocols.

To define the clusters, we completed the following steps:

1. Clusters of each area were extracted from a 2mm standard template using Harvard-Oxford and Juelich atlases.
2. If applicable, clusters from both atlases were added up.
3. Clusters were then thresholded and binarized.
4. Binarized and thresholded clusters were applied to the group-level thresholded z statistical maps for each preprocessing protocol to create a functional mask for each protocol.
5. Individual averaged parameter estimates were extracted within area by performing featquery on the individual second-level copes and varcopes using the functional masks defined in 4.

Resulting averaged parameter estimates were compared by performing a three-way repeated-measures ANOVA [maps: 2 levels (cope/varcope) x brain areas: 5 levels (ACC, S1, insula, white matter, CSF) x preprocessing protocols: 3 levels (FIX, CompCor, standard)] after extreme outliers were excluded from the analyses. Extreme outliers were defined as individual averaged parameter estimates that were greater than the third quartile + 3*interquartile range (Q3 + 3*(Q3-Q1)) or that were smaller than the first quartile – 3*interquartile range (Q1 – 3*(Q3-Q1)). Post-hoc analyses were performed when appropriate. Bonferroni correction for multiple comparisons was applied when appropriate. A significance p threshold was defined at 0.05. These analyses were performed using Rstudio version 1.3.1093.

## Results

### Differences in brain activation associated with high intensity heat stimulation using a block design between preprocessing protocols

Group-level GLM analyses were performed to assess the BOLD signals associated with high intensity heat stimulation after each preprocessing protocol, i.e. standard, with FIX noise-reduction technique, and with CompCor noise-reduction technique.

Data preprocessed with all three preprocessing protocols revealed changes in activations in similar areas (Figure 2). Increased activation was observed in grey matter areas encompassing the anterior cingulate cortex (ACC), the primary sensorimotor cortex (S1M1), the insula, the basal ganglia, and the thalamus. Deactivation was observed primarily in grey matter areas including the amygdala and hippocampus, as well as in the posterior cingulate cortex (PCC) and precuneus.

**Figure 2.**
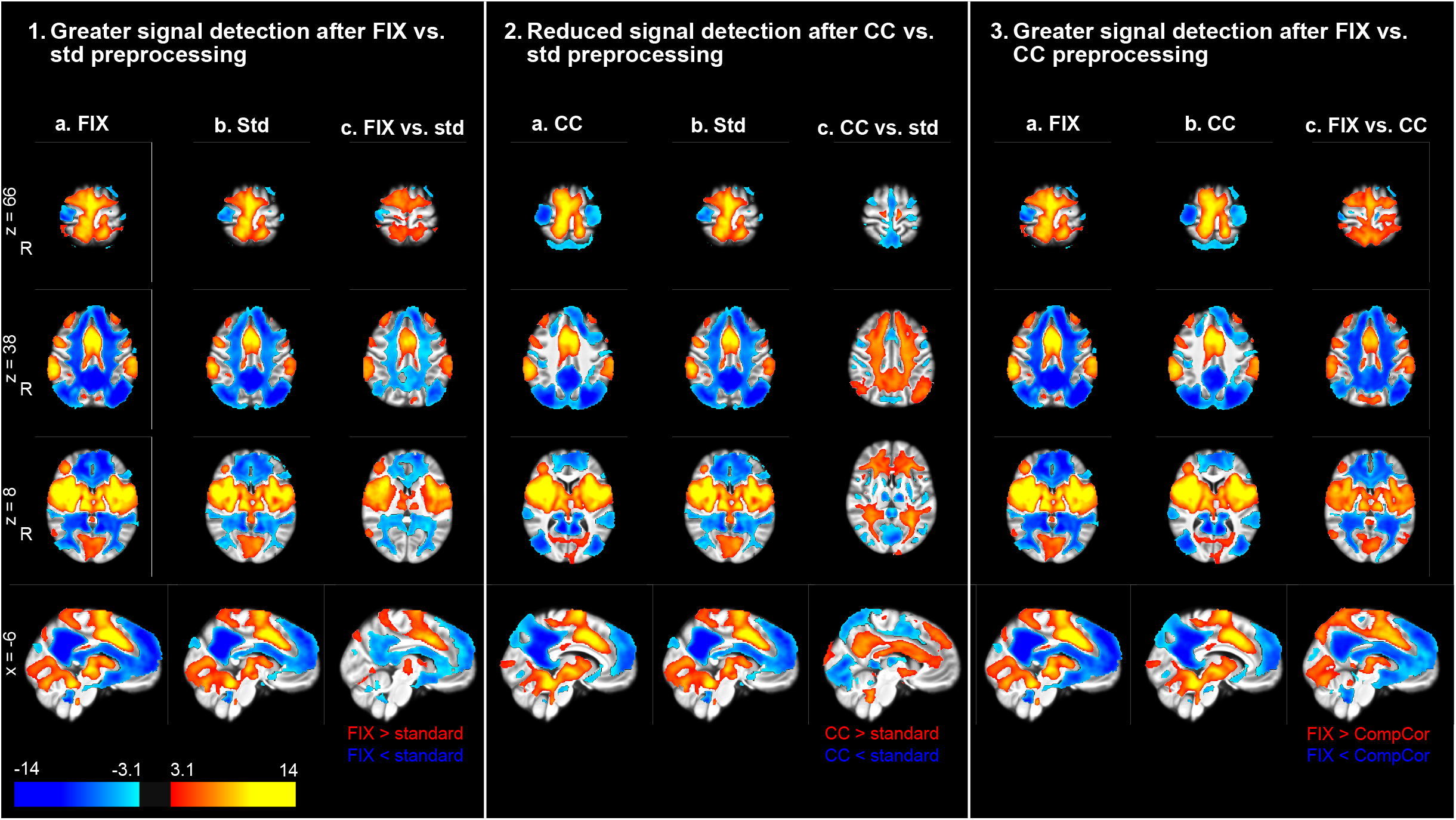
Brain activation associated with high intensity heat stimulus after preprocessing with FIX noise-reduction technique is greater than after preprocessing with CompCor noise-reduction technique and standard preprocessing. (1.a) Following FIX preprocessing, noxious heat stimuli are associated with increased and decreased brain activation in grey matter areas previously associated with pain and in the white matter; (1.b) brain activation associated with noxious heat stimuli after standard (std) preprocessing is observed in similar areas as after FIX preprocessing; (1.c) The comparison of brain activation after preprocessing protocol with FIX noise-reduction technique to brain activation after standard preprocessing shows greater positive and negative signal detection after FIX in grey matter areas previously associated with pain and in the white matter. (2.a) Increased and decreased brain activation associated with noxious heat stimuli after CompCor (CC) preprocessing is observed in grey matter areas previously associated with pain; (2.b) Increased and decreased brain activation associated with noxious heat stimuli after standard preprocessing was shown in grey matter areas previously associated with pain and in the white matter; (2.c) In contrast with the FIX vs. std comparison, signal appeared to be removed by CompCor. Greater positive and negative signal in areas previously associated with pain is detected after standard preprocessing compared to CompCor preprocessing. (3.a) Brain activation associated with noxious heat stimuli after FIX preprocessing is observed in the same areas as described in 1.a; (3.b) Brain activation associated with noxious heat stimuli after CC preprocessing as described in 2.a; (3.c) Consistent with the patterns of activation noted in the comparisons with std preprocessing, the direct comparison of brain activation after FIX preprocessing and brain activation after CC preprocessing shows greater positive and negative signal detection in grey matter areas previously associated with pain after FIX preprocessing.

Deactivation in the white matter was also observed following standard and FIX preprocessing. Application of the FIX noise-reduction technique enhanced the magnitude of meaningful signals of interest compared to the standard preprocessing protocol (Figure 2.1). The grey matter regions mentioned above and typically activated during high intensity heat stimulation exhibited greater activation with FIX preprocessing than standard preprocessing. Conversely, the grey matter regions listed above and typically deactivated during high intensity heat stimulation exhibited greater deactivation with FIX vs. standard preprocessing. Similarly, reduced BOLD signals in the white matter exhibited greater reductions with FIX vs. standard preprocessing.

The comparison of brain activation associated with high intensity heat stimulus after CompCor preprocessing and after standard preprocessing yielded a loss in the detection of signals of interest after the CompCor preprocessing protocol (Figure 2.2). Compared to standard preprocessing, smaller activations and deactivations in grey matter areas typically associated with noxious heat stimulation were found after CompCor preprocessing. Similarly, smaller deactivation was detected in the white matter following CompCor preprocessing than following standard preprocessing.

Finally, a T-test comparing brain activation associated with high-intensity heat stimulation after FIX preprocessing and brain activation after CompCor preprocessing showed increased signal detection after the FIX preprocessing vs. after CompCor preprocessing (Figure 2.3). Greater increased and decreased activations were found in grey matter areas typically associated with noxious heat stimulation following FIX preprocessing vs. CompCor preprocessing. Deactivation in the white matter was greater after FIX preprocessing vs. CompCor preprocessing.

**Figure 3.**
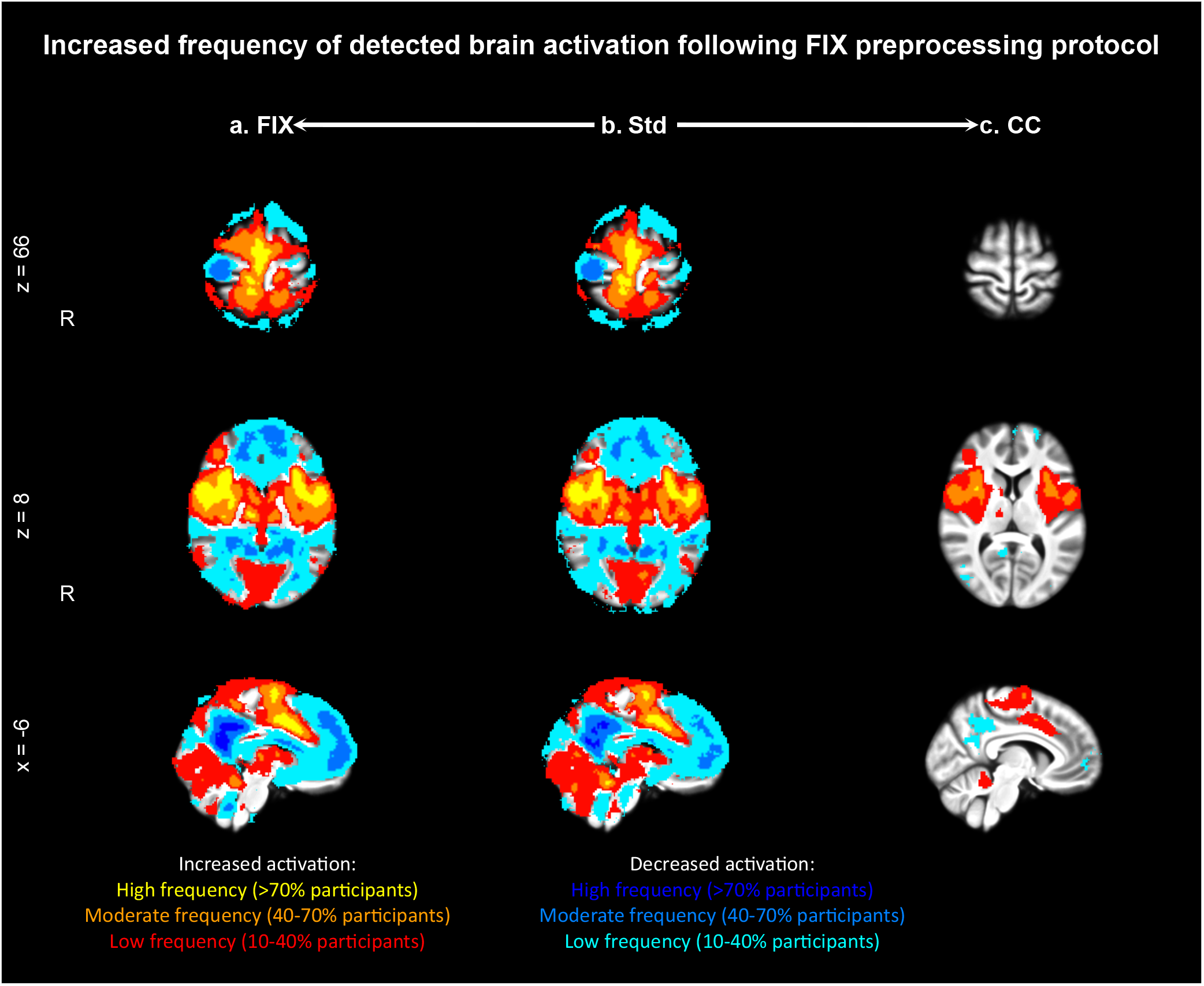
Frequency of activation, i.e. percentage of participants with similar brain activation, is greater after FIX preprocessing compared to the other two preprocessing protocols. Frequency of increased activation is represented in yellow for high frequency (activation detected in more than 70% of participants), in orange for moderate frequency (40-70% of participants), and in red for low frequency (10-40% of participants). Frequency of decreased activation is represented in dark blue for high frequency, in blue for moderate frequency, and in cyan for low frequency. a) Frequency of increased activation after FIX preprocessing was high in areas including S1M1, the ACC, and the insula, moderate around these areas and low around the moderate areas. Decreased activation was observed with high frequency in areas including the precuneus and PCC, with moderate frequency around these areas and in the white matter, and with low frequency in areas surrounding the moderate-frequency areas; b) Compared to frequency following FIX preprocessing, high, moderate, and low frequency of increased and decreased activation after standard (std) preprocessing was observed in similar but smaller areas; c) In contrast to frequency observed after FIX and std preprocessings, frequency of increased and decreased activations was observed at lower levels and in smaller areas. High frequency of increased activation was only observed in a small area of the insula, while moderate- and low-frequency activations were observed in the insula and ACC. Decreased activation was only observed in low frequency in the precuneus and PCC.

### Frequency of activation across participants

Frequency analyses were performed to assess the percentage of participants exhibiting brain activation in similar areas in response to high intensity heat stimulation following each preprocessing protocol.

These analyses revealed that the maximum percentage of participants showing brain activation and deactivation in similar areas was greater following FIX and standard preprocessing than following CompCor preprocessing (Table 2).

**Table 2.** Maximum percentage of participants showing similar brain activation in response to highly noxious stimulus is greater following preprocessing with FIX or standard preprocessing than following preprocessing with CompCor. Increased activation represents the maximum percentage for areas with increased activation, while decreased activation shows the maximum percentage for areas with decreased activation.

| **Preprocessing pipeline** | **Increased activation** | **Decreased activation** |
| --- | --- | --- |
| **FIX** | 94% | 79% |
| **Standard** | 97% | 84% |
| **CompCor** | 71% | 39% |

High frequency of activation and deactivation, i.e. activation/deactivation detected in more than 70% of participants, was detected differently depending on the preprocessing protocol (Figure 3; yellow/dark blue activation). Following CompCor preprocessing, increased activation was only detected in the right insula and no deactivation was detected at that frequency. Following FIX and standard preprocessing, increased activation in the right S1M1, the bilateral ACC, and the bilateral insula was detected at that frequency. Deactivations following FIX and standard processing were detected in the bilateral precuneus and PCC at the same frequency. Moderate frequency of increased activation (Figure 3; orange activation), i.e. activation that was detected in 40 to 70% of the participants, was detected in areas surrounding the areas detected with high frequency and was consistently greater in terms of spatial extent and frequency following FIX preprocessing vs. following standard and CompCor preprocessing. Moderate frequency of deactivation (Figure 3; blue deactivation) was detected in areas surrounding the ones detected with high frequency and in white matter areas. The spatial extent and frequency of deactivations detected at that frequency were greater following FIX preprocessing vs. standard preprocessing.

Finally, low frequency of activation and deactivation (Figure 3; red/cyan activation), which was detected in 10 to 40% of the participants, was detected following all our preprocessing protocols. This frequency was consistently greater in terms of spatial extent and frequencies following FIX preprocessing vs. CompCor and standard preprocessing and following CompCor preprocessing vs. standard preprocessing. Following FIX and standard preprocessing, low-frequency activations and deactivations were detected in areas surrounding the areas activated at moderate frequency. Following CompCor preprocessing, increased activation was detected at a low frequency in areas surrounding the areas activated at a moderate frequency. Deactivation following CompCor preprocessing was only detected at a low frequency and only in a small area of the precuneus and PCC.

### Differences in brain activation associated with a single high intensity heat stimulus using a single-epoch design between preprocessing pipelines

GLM analyses were performed to investigate the effect of each preprocessing protocol on brain activation associated with a single epoch, i.e. a single high intensity heat stimulus. Results from these analyses (Figure 4) showed increased brain activation after FIX, CompCor, and standard preprocessing in areas such as ACC, S1M1, the insula, basal ganglia, and the thalamus.

**Figure 4.**
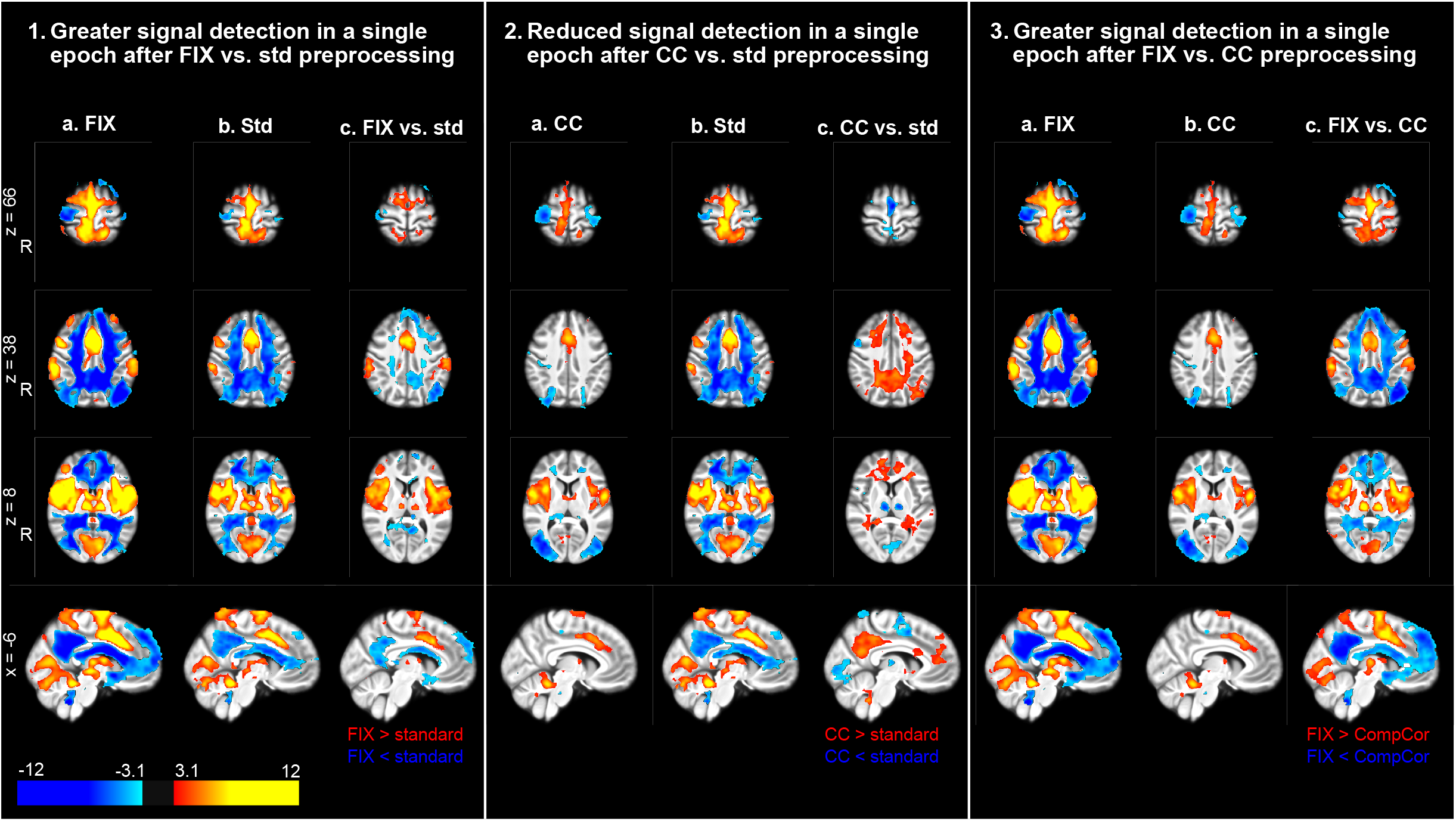
Signal detection associated with a single high intensity heat stimulus after FIX preprocessing is greater than after CompCor and standard preprocessing. (1.a) Increased and decreased brain activation associated with a single noxious heat stimulus after FIX preprocessing is observed in grey matter areas consistent with pain processing and in the white matter; (1.b) Increased and decreased brain activation associated with a single noxious heat stimulus after standard (std) preprocessing was observed in areas similar to 1.a; (1.c) The comparison of brain activation after FIX preprocessing and brain activation after standard preprocessing shows greater positive and negative signal detection after FIX preprocessing. (2.a) Increased and decreased brain activation associated with a single noxious heat stimulus after CompCor (CC) preprocessing was observed in grey matter areas consistent with pain processing; (2.b) Brain activation associated with a single noxious heat stimulus after standard preprocessing is identical to the one described in 1.b; (2.c) Compared to standard preprocessing, signal appears to be removed by the CC preprocessing. (3.a) Brain activation associated with a single noxious heat stimulus after FIX preprocessing is identical to the one described in 1.a; (3.b) Brain activation associated with noxious heat stimulus after CC preprocessing as described in 2.a; (3.c) Similarly to the comparison between CC and std preprocessing, signals appear to be removed by CC preprocessing compared to FIX preprocessiong. Greater positive and negative signals is detected following FIX preprocessing.

Deactivation was observed in the precuneus and PCC after preprocessing with FIX noise-reduction technique and after standard preprocessing, but not after preprocessing with CompCor noise-reduction technique.

After FIX preprocessing compared to standard preprocessing, an increased signal was detected at a single stimulus level (Figure 4.1): greater activation was found in grey matter areas previously described and typically associated with noxious stimulation. Deactivation in grey matter areas, including the PCC and precuneus, and in the white matter was better detected following FIX preprocessing.

When comparing the preprocessing protocol including CompCor noise-reduction technique with the standard preprocessing protocol at a stimulus level, better signal detection can be observed after standard preprocessing (Figure 4.2). Increased activation was detected after standard preprocessing in grey matter areas typically associated with noxious stimulation. Deactivation was also better detected after standard preprocessing in grey matter areas, including the PCC and the precuneus and the white matter.

Signal detection increased greatly after preprocessing including the FIX noise-reduction technique compared to preprocessing with CompCor noise-reduction technique at a single stimulus level (Figure 4.3). Increased activation was detected following preprocessing with FIX in grey matter areas associated with noxious stimulation. Deactivation was better detected after preprocessing with FIX in grey matter areas, including the PCC and the precuneus and the white matter.

### Noise and signal detection after each preprocessing pipeline

A three-way repeated-measures ANOVA was performed to investigate the interactive effect between maps (cope/varcope), brain areas, and preprocessing protocols on averaged parameter estimates. Results of this ANOVA showed a significant interaction between maps, brain areas, and preprocessing protocols (F(3.08,209.44) = 55.564, p < 0.0001).

Given this significant result, two two-way repeated-measures ANOVA were performed to compare averaged parameter estimates between brain areas and preprocessing protocols within map. Results of these analyses showed significant interactions between brain areas within map and preprocessing protocols after Bonferroni correction (cope: F(2.97,285) = 53.7, p < 0.0001; varcope: F(1.26, 85.5) = 31.8, p < 0.0001).

To further investigate these interactions, one-way ANOVAs were performed to establish the effect of preprocessing protocols within map and brain areas. Results after Bonferroni correction showed a significant main effect of the preprocessing protocols on averaged parameter estimates in the cope map in the following areas: ACC (F(1.4,139) = 123, p < 0.0001), insula (F(1.79,179) = 129, p <0.0001), S1 (F(1.74,172) = 117, p < 0.0001), and white matter (F(1.55,153) = 65.7, p < 0.001). A significant main effect of the preprocessing protocols was also found in the varcope map in all the brain areas studies (ACC: F(1.12,106) = 180, p < 0.0001; insula: F(1.15,112) = 285, p < 0.0001; S1: F(1.45, 139) = 117, p < 0.0001; white matter: F(1.1,102) = 162, p < 0.0001; CSF: F(1.25, 98.4) = 39.6, p < 0.0001).

Finally, pairwise paired T-tests were performed within map and brain area to identify differences in parameter estimates between preprocessing protocols based on the significant main effects of the preprocessing protocols identified in the one-way ANOVAs. In the cope image, Bonferroni-corrected results (Figure 5, left panel) showed significant differences in the ACC between all protocols (FIX – CompCor: t(99) = -13.29, p < 0.0001; FIX – standard: t(99) = - 11.49, p < 0.0001; CompCor – standard: t(99) = -3.62, p = 0.001). In the insula, significant differences were found between FIX and CompCor preprocessing (t(100) = -15.67, p < 0.0001) and between CompCor and standard preprocessing (t(100) = -13.51, p < 0.0001). Significant differences were found between all the preprocessing protocols in S1 (FIX – CompCor: t(99) = - 8.8, p < 0.0001; FIX – standard: t(99) = -4.71, p < 0.0001; CompCor – standard: t(99) = -10.65, p < 0.0001). Finally, significant differences in the white matter were observed between FIX and CompCor preprocessing (t(99) = 10.81, p < 0.0001) and between CompCor and standard preprocessing (t(99) = 8.5, p < 0.0001).

**Figure 5.**
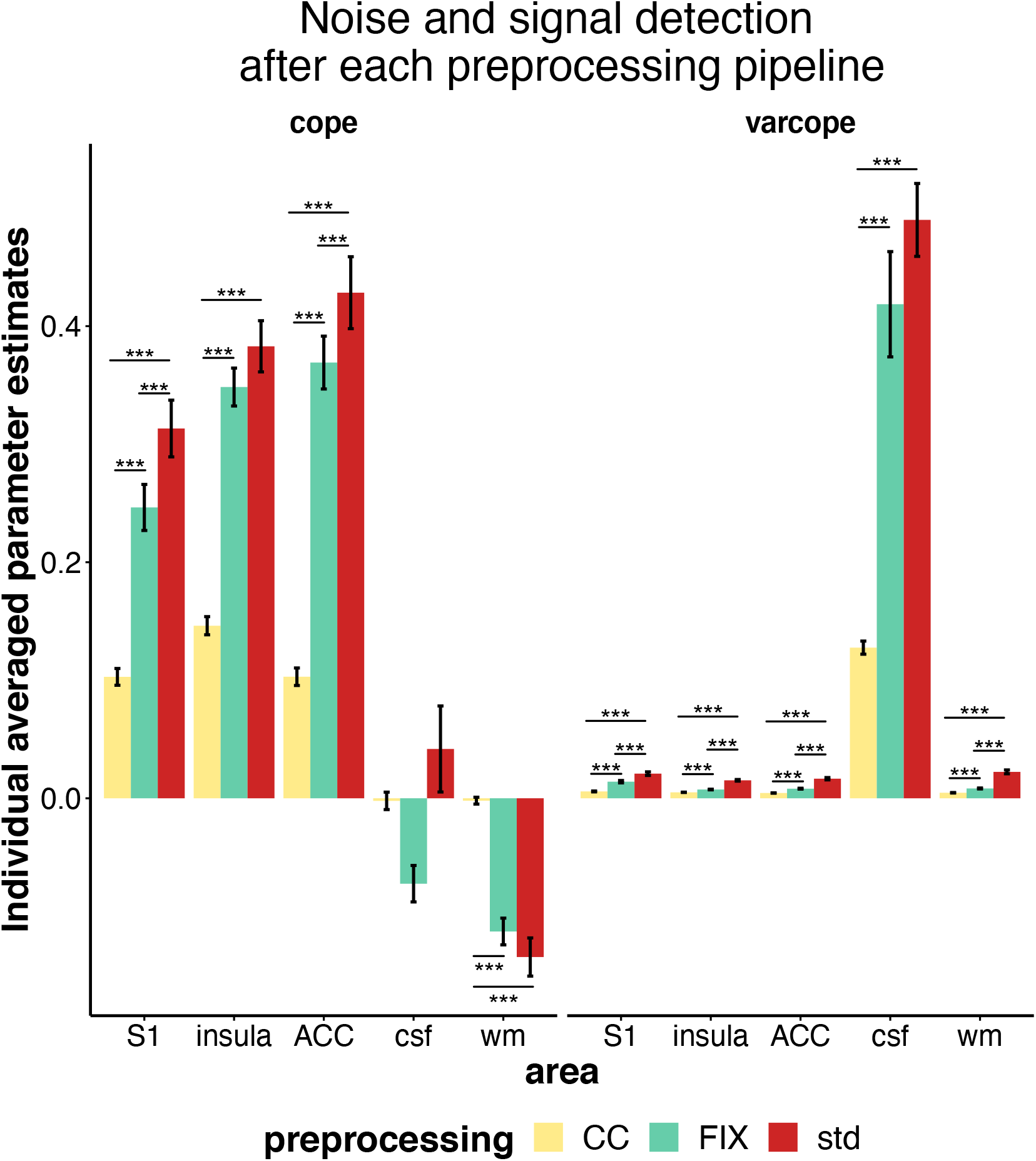
FIX conserves significantly more signal individually in areas of interest than CompCor, while removing more noise than standard preprocessing. Individual averaged parameter estimates in cope (contrast parameter estimates; left panel) and varcope (variance of contrast parameter estimates; right panel) images are represented for each preprocessing pipeline (CompCor (CC): yellow, FIX: green; standard (std): red) for the following areas of interest: ACC: anterior cingulate cortex; csf: cerebrospinal fluid; insula; S1: primary somatosensory cortex; wm: white matter. Signal detection was significantly greater after standard preprocessing than after FIX preprocessing in S1 and ACC and than after CompCor in S1, insula, ACC, and white matter. Similarly, significantly greater signal detection following FIX preprocessing than after CompCor preprocessing was observed in the same areas. Remaining noise in the images was always significantly greater following standard preprocessing and FIX preprocessing than following CompCor preprocessing in all areas investigated. Remaining noise was also significantly greater in S1, insula, ACC, and white matter following standard preprocessing than following FIX preprocessing. *** p < 0.0001.

In the varcope image (Figure 5, right panel), significant differences were found between all preprocessing protocols in the ACC (FIX – CompCor: t(94) = -14.3, p < 0.0001; FIX – standard: t(94) = -11.77, p < 0.0001; CompCor – standard: t(94) = -14.37, p < 0.0001), in the insula (FIX – CompCor: t(97) = -14.07, p < 0.0001; FIX – standard: t(97) = -15.27, p < 0.0001; CompCor – standard: t(97) = -18.34, p < 0.0001), in S1 (FIX – CompCor: t(96) = -10.84, p < 0.0001; FIX – standard: t(96) = -7.83, p < 0.0001; CompCor – standard: t(96) = -12.13, p < 0.0001), and in the white matter (FIX – CompCor: t(93) = -11.37, p < 0.0001; FIX – standard: t(93) = -11.49, p < 0.0001; CompCor – standard: t(93) = -13.86, p < 0.0001). In the CSF, significant differences were found between FIX and CompCor preprocessing (t(77) = -6.01, p < 0.0001) and between CompCor and standard preprocessing (t(77) = -16.47, p < 0.0001).

Overall, the FIX preprocessing protocol preserved a lot more signal of interest at an individual level than the CompCor preprocessing protocol while removing less noise (Figure 5). Comparing the standard preprocessing protocol to the CompCor preprocessing protocol showed similar results. Finally, the standard preprocessing protocol preserved slightly more signal of interest at an individual level but also removed less noise than the FIX preprocessing protocol.

### Differences in brain activation associated with high intensity auditory stimulation using a block design between preprocessing protocols

Non-noxious auditory stimulation was used as a control modality to investigate the effect of the different preprocessing protocols on cerebral responses to stimulation that has no strong associated physiological response.

Analyses of brain activations associated with high-intensity auditory stimulation following standard, FIX, and CompCor preprocessing protocols showed increased activations in areas previously associated with auditory perception, including the primary auditory cortex (A1), ACC, and the insula (Figure 6). Deactivations were seen after all our preprocessing protocols in grey matter areas including S1M1, and in the white matter.

**Figure 6.**
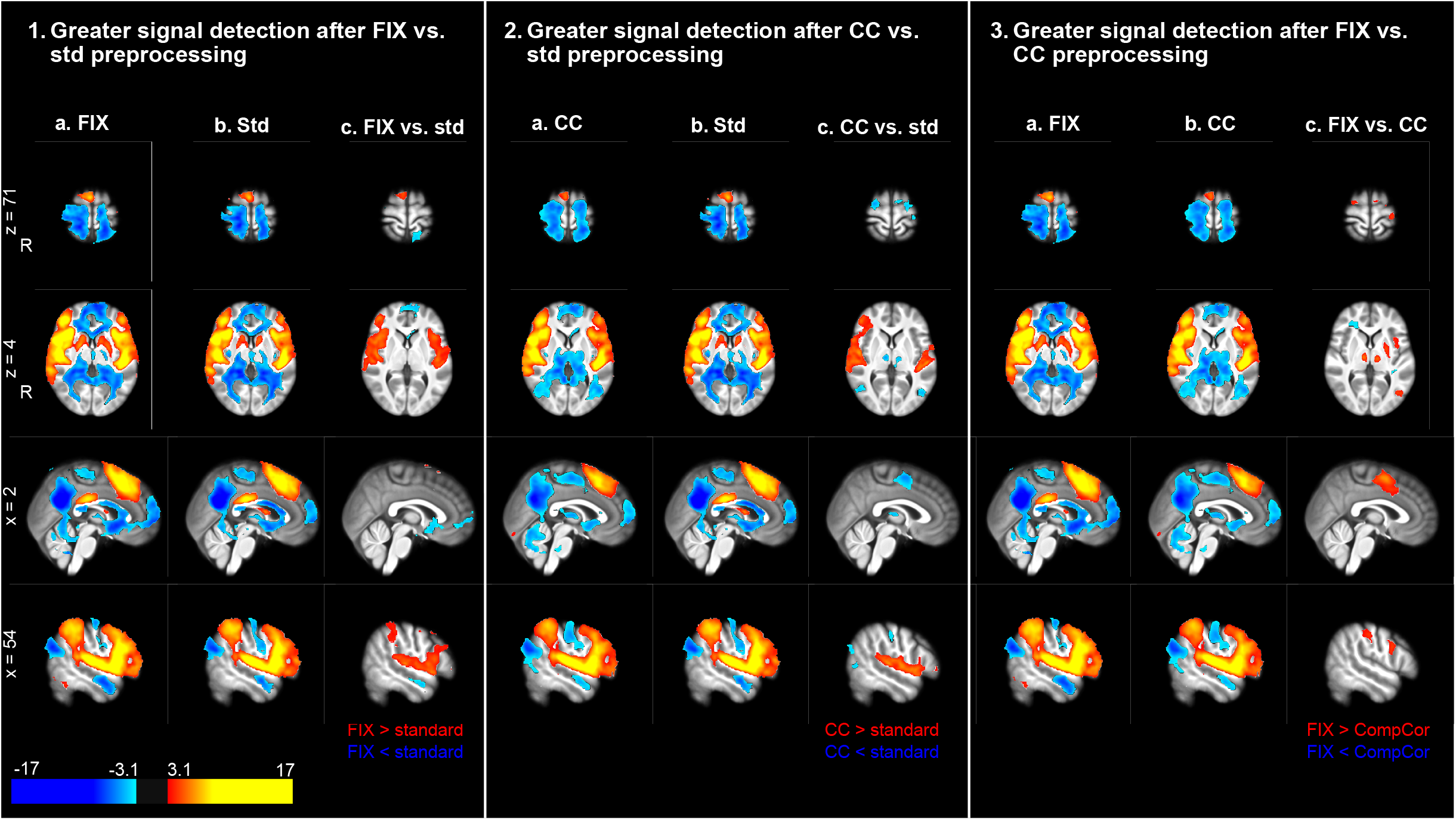
Brain activation associated with high intensity auditory stimulus after FIX preprocessing is greater than after CompCor and standard preprocessing. (1.a) Increased brain activation associated with auditory stimuli after FIX preprocessing is observed in areas including A1, the insula, putamen, and ACC. Decreased brain activation was observed primarily in precuneus and PCC, as well as in the white matter; (1.b) Increased and decreased brain activation associated with auditory stimuli after standard (std) preprocessing was similar to changes in activation observed after FIX preprocessing; (1.c) The comparison of brain activation after FIX preprocessing and brain activation after standard preprocessing showed positive and negative greater signal detection after FIX preprocessing in similar areas. (2.a) Increased brain activation associated with auditory stimuli after CompCor (CC) preprocessing was observed in similar brain areas as after FIX and standard preprocessing, while decreased activation was observed primarily in PCC and precuneus; (2.b) Brain activation associated with auditory stimuli after std preprocessing is identical to changes described in 1.b; (2.c) Similarly to the comparison between FIX and std preprocessings, brain activation after CompCor preprocessing showed greater positive and negative signal detection after CompCor in A1 and thalamus than after std preprocessing. (3.a) Brain activation associated with auditory stimuli after FIX preprocessing is identical to the changes described in 1.a; (3.b) Brain activation associated with auditory stimuli after CC preprocessing is identical to changes described in 2.a; (3.c) Similarly to the comparison of brain activation after FIX vs. std preprocessing, brain activation after FIX preprocessing was greater than after CC preprocessing in similar areas.

After FIX preprocessing compared to standard preprocessing, the detection of the signal of interest was increased in grey matter areas, including A1 and the insula (Figure 6.1). Similarly, improved signal detection in A1 was observed when the CompCor noise-reduction technique was applied compared to the standard preprocessing protocol (Figure 6.2). In addition, following CompCor preprocessing, greater deactivation was evident in the left S1M1 and the thalamus compared to standard preprocessing. In contrast, decreased detection of activation in the ACC was found following CompCor preprocessing vs. standard preprocessing.

Finally, the comparison of FIX preprocessing with CompCor preprocessing yielded three main findings: (1) increased signal detection was observed in grey matter areas showing positive activation, including ACC and putamen; (2) greater deactivation in the thalamus was found after CompCor preprocessing; (3) small areas of the white matter were more deactivated following preprocessing with FIX.

## Discussion

FIX and CompCor preprocessing protocols reduced noise in task-related fMRI data. Specifically, both noise reduction techniques enhanced the ability to detect activation during an auditory stimulus. However, for tasks that elicit correlated white matter responses, such as noxious heat stimulation, FIX increases signal detection while CompCor removes more signals of interest. FIX preprocessed data displayed robust activation in this task, while considerably less activation was detected with CompCor, likely due to correlated white matter responses. These findings underscore the benefits of data cleaning protocols on task-related data, while emphasizing the fact that the wrong choice of a data cleaning protocol can have a deleterious impact on the ability to assess brain activation.

At a group level, analyses based on a block design showed that greater brain activation associated with high intensity heat stimulation was detected after FIX preprocessing than after CompCor or standard preprocessing. Following FIX preprocessing, increased activation was detected in grey matter regions typically associated with noxious heat stimuli, including the ACC, the insula, and S1. Increased deactivation was detected in grey matter areas that are part of the default mode network and typically decreased in response to painful stimuli, including the precuneus and PCC.

The superior performance of the FIX noise-reduction technique is further supported by the analyses performed on a single high intensity heat stimulus using a single epoch design. Single epoch designs are more sensitive to noise than the block design approaches used in the other analyses (Koyama et al., 2003). Greater activations and deactivations were detected in grey matter areas similar to those seen in the block design following FIX preprocessing in comparison with activations/deactivations detected after CompCor and standard preprocessing. This clearly highlights the capacity of the FIX noise-reduction technique to increase the signal-to-noise ratio during the cleaning process.

To understand the impact of different noise-reduction techniques at the level of the single participant, we examined how many individuals activated specific brain regions in a suprathreshold fashion (i.e. frequency of activation). Results from this block-design based frequency analysis showed: (1) greater frequency of detection and spatial extent within grey matter areas associated with response to noxious stimulation following FIX preprocessing compared to standard preprocessing; (2) the maximum percentage of participants showing similar within-voxel activation across the whole brain, as well as the frequency of detection and spatial extent within grey matter areas usually associated with noxious stimulation was lower following CompCor preprocessing compared to standard preprocessing; (3) when comparing the impact of the FIX preprocessing and CompCor preprocessing, the maximum percentage of participants, as well as the frequency of detection and the spatial extent within grey matter areas typically associated with response to noxious stimulation were greater following FIX preprocessing. Thus, the benefits of FIX noise reduction are evident at the individual level.

The analysis of the individual averaged parameter estimates in areas of interest including the ACC, insula, and S1, showed that FIX and standard preprocessing conserved more signal than CompCor preprocessing, while both FIX and CompCor preprocessing removed more noise than standard preprocessing. Similarly, FIX and standard preprocessing conserved more signal in the white matter than CompCor preprocessing, while FIX and CompCor preprocessing removed more noise. Finally, FIX and CompCor preprocessing removed more noise than standard preprocessing in the cerebrospinal fluid, while there was no significant difference in the conserved signal between the three preprocessing protocols.

While the CompCor preprocessing protocol removed the most noise at individual and group levels, it also removed signals of interest. This was shown by fewer activated areas and a lower magnitude of activation in our GLM analyses and by detecting similar patterns of brain activation in only a low percentage of participants. This is likely due to some signal of interest being temporally associated with white matter response in the noxious heat task. CompCor noise-reduction technique relies on the assumptions that all the signal present in the white matter and CSF is noise and that noise in the grey matter is temporally correlated with noise in the white matter and CSF. Given that our noxious heat task is associated with significant white matter response, by regressing out the signals detected in this area, temporally related signal of interest in the grey matter was removed as well. Unlike the common beliefs that responses in the white matter are the result of noise, our results suggest that in tasks including noxious stimulation, changes in white matter signal might be representative of some systemic response to the stimulus in line with recent publications (Grajauskas et al., 2019; Li et al., 2020). Unfortunately, the design of our study does not allow us to define the source(s) of this response. However, global changes in cerebral blood flow may potentially contribute to the white matter signal changes, as such changes have been previously identified during intense noxious stimuli (Coghill et al., 1998). Further studies are needed to delineate the mechanisms underlying these transient changes in white matter signal intensity.

Interestingly, the effect of the preprocessing protocol, and in particular of the denoising technique, seems to lessen in tasks with less associated white matter response, as shown in the analyses of our control condition. The analyses performed on our auditory paradigm showed that, although FIX preprocessing still increases signal detection compared to standard and CompCor preprocessing, the difference in terms of signal of interest can be minimal (see the comparison between FIX and CompCor preprocessing, Figure 6). This suggests that the impact of the preprocessing protocols is lesser in tasks that are not associated with significant white matter response, such as our auditory task, compared to our noxious heat paradigm. These results clearly show that different preprocessing protocols impact signal-to-noise ratio differently depending on the tasks. Furthermore, these results suggest that, among the preprocessing protocols tested in this study, the inclusion of the FIX noise-reduction technique in the preprocessing protocol is the most appropriate data cleaning approach for task fMRI data associated with an important white matter response and are interested in individual responses.

In conclusion, preprocessing fMRI data is always a delicate balance between conserving the signal of interest and removing the noise. The choice of preprocessing protocols can significantly influence the signal-to-noise ratio. The efficacy of these noise reduction techniques can vary dynamically according to the nature of the signals within the brain. Therefore, studies should carefully consider the choice of the preprocessing protocols based on the type of tasks that will be used. Our results suggest that in tasks with important associated white matter responses and tasks without such responses, a preprocessing protocol including a FIX noise-reduction technique might be a good solution to achieve this balance.

## Notes

### Competing Interest Statement

The authors have declared no competing interest.

